# Cancer cell-autonomous cGAS-STING response confers drug resistance

**DOI:** 10.1101/2021.10.23.465546

**Authors:** Qian-Ming Lv, Shi-Yi Wang, Hui-Min Lei, Ke-Ren Zhang, Ya-Bin Tang, Ying Shen, Li-Ming Lu, Hong-Zhuan Chen, Liang Zhu

## Abstract

As an evolutionarily conserved DNA-sensing machinery in innate immunity, the cGAS-STING pathway has been reported to play an important role in immune surveillance and tumor suppression. Recent evidence suggests an intriguing tumor- and metastasis-promoting effect of this signaling pathway, either in a cancer cell-autonomous or a cancer cell-nonautonomous, bystander cell-mediated manner. Here, we show a new face of cGAS-STING signaling whose activation in a cancer-cell-autonomous response manner confers drug resistance. Targeted or conventional chemotherapy drug treatment induced cancer cell cytosolic DNA accumulation and triggered subsequent cGAS-STING signaling activation in cancer cell lines and the human cell-derived xenograft tumors. This activation promoted an acquisition and maintenance of drug resistance which was prevented and overcome in vitro and in vivo by blockade of STING signaling. This finding highlights a new face of cGAS-STING signaling and an ability of cancer cells to hijack the evolutionarily conserved inflammatory signaling to counteract drug stress and warrants a caution in combining STING agonist with targeted or conventional chemotherapy drug treatment, a strategy prevailing in current clinical trials.

**Statement of significance:** cGAS-STING signaling has long been recognized as playing a key role in triggering antitumor immunity. We reveal a new face of cGAS-STING signaling and an ability of cancer cells autonomously to hijack the evolutionarily conserved inflammatory signaling to counteract drug stress.

## Introduction

STING (stimulator of interferon genes) serves as a linchpin in the cytosolic DNA-sensing pathway. Activated by cyclic GMP-AMP (cGAMP) that is produced by cyclic GMP-AMP synthase (cGAS) sensing cytosolic double-stranded DNA (dsDNA), STING boosts downstream inflammatory signaling (1,2). Discovered primarily as an important machinery in innate immunity and host defense against microbial pathogens, cGAS-STING has been extendedly depicted as playing a key role in triggering antitumor immunity (1,2).

The putative tumor-suppressive effect of STING motivates the research and development of STING agonists for cancer immunotherapy. However, the limited efficacy of these STING agonist in clinical trials warrants a more comprehensive understanding of cGAS-STING characteristics (1,2). Recently, evidence suggests an intriguing cancer and metastasis-promoting effect of cancer cell-intrinsic cGAS-cGAMP-STING activation, either in a cancer cell-autonomous response manner (3,4) or in a cancer cell-nonautonomous, bystander cell-mediated manner (5), indicating the many faces of cGAS-STING signaling in cancer biology other than triggering antitumor immunity which has received more attention (1,2,6,7).

Resistance is one of the biggest challenges for cancer treatment (8). Based on our in vitro and in vivo cancer targeted therapy resistance models (9–11), we repeatedly and curiously noticed that an activated rather than suppressed status was achieved in cGAS-STING signaling when these cancer cells acquired resistance to therapeutic drugs. We hypothesized that the cells that can upregulate cGAS-STING signaling would acquire the ability to cope with drug stress and evolve drug resistance.

Herein, we show that a cancer-cell-autonomous response of the cell-intrinsic cGAS-STING activation triggered by targeted or chemotherapy drugs confers drug resistance. This finding highlights a novel face of cGAS-STING signaling and a skill of cancer cells hijacking the evolutionarily conserved inflammatory signaling to counteract drug stress and warrant a caution in combining a STING agonist with anti-cancer drugs, a prevailing strategy in current clinical trial designs.

## Results

### Hyperactivation of cGAS-STING signaling in drug-resistant tumor and tumor cells

Transcriptomic analysis of the differentially expressed genes between the various lines of EGFR TKI-resistant cells (Supplementary Fig. S1A, top panel) versus their isogenic parental sensitive cells demonstrated an enrichment for cytosolic DNA sensing pathway (Supplementary Fig. S1B, top panel). The enrichment was recapitulated in chemotherapy drug cisplatin-resistant cells (Supplementary Fig. S1B, bottom panel) and in vivo drug-resistant models (Supplementary Fig. S1A, bottom panel) where the erlotinib treatment-relapsed tumors showed a hyperactivation of cytosolic DNA sensing signature compared with the sensitive tumors (Fig. 1A). Likewise, cGAS-STING downstream target genes were upregulated in resistant cells (Fig. 1B, Supplementary Fig. S1C).

**Figure 1.**
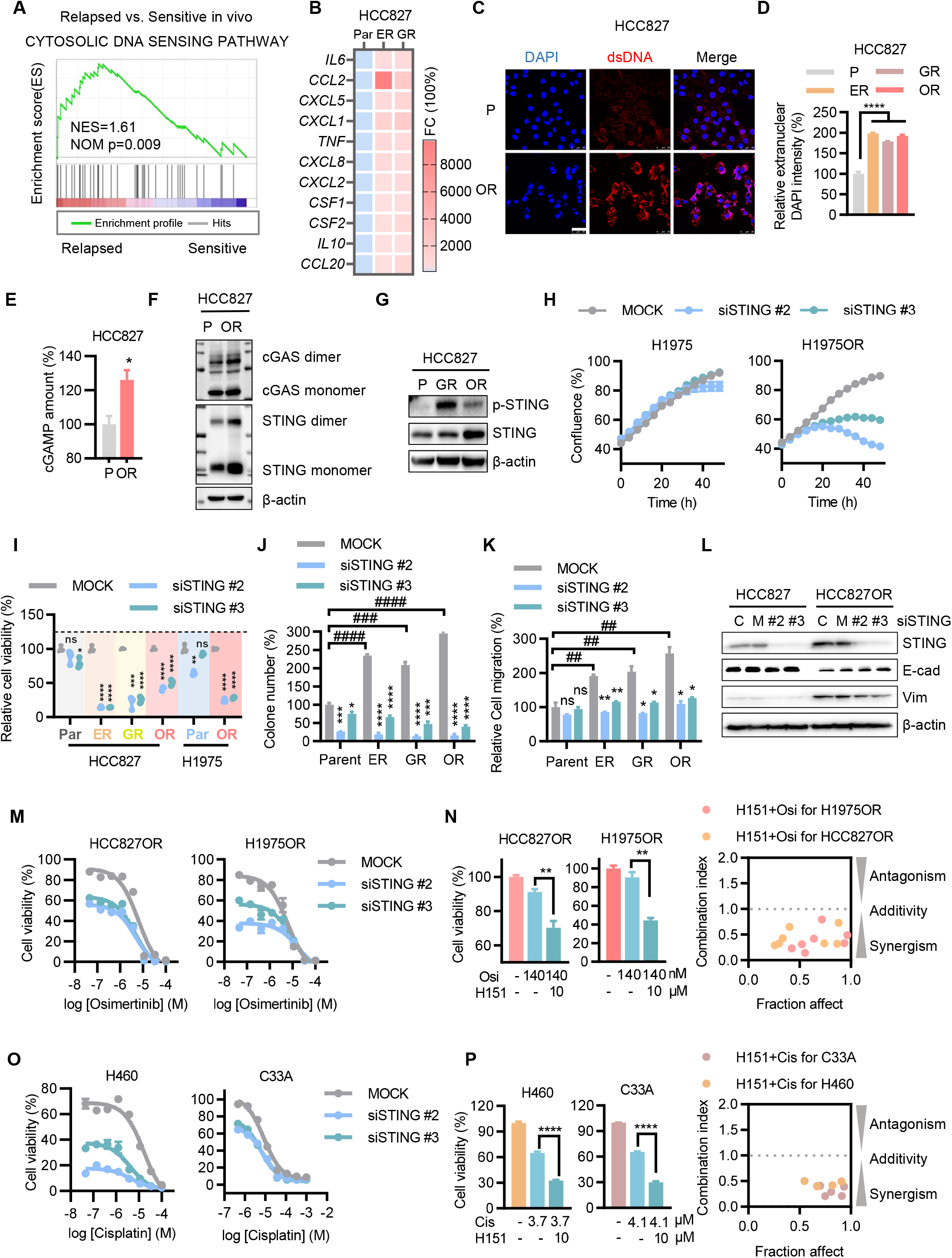
Drug-resistant cancer cells demonstrate and depend on an activated cGAS-STING signaling. (A) Gene set enrichment analysis (GSEA) of RNA-seq data indicates a hyperactivation of cytosolic DNA sensing pathway in erlotinib treatment-relapsed tumors (n=5) compared with the sensitive tumors (n=6). The schematic diagram of the establishment of the in vivo models is shown in Supplementary Fig. S1A. (B) Heatmap shows expression difference of cGAS-STING signaling downstream genes between HCC827 TKI-resistant (ER and GR) and parental (Par) cells assayed by RNA-seq analysis. ER and GR represent erlotinib and gefitinib resistance, respectively. (C) Double-strand DNA (dsDNA) accumulation in the cytoplasm of HCC827OR resistant versus parental (P) cells assayed by anti-dsDNA antibody (1:50, Abcam)-based immunofluorescence staining analysis. Scale bar: 50 μm. (D) Extranuclear DNA accumulation in HCC827 resistant (ER, GR, OR) versus parental (P) cells assayed by quantification of relative extranuclear DAPI fluorescence intensity. (E) Accumulation of intracellular cGAMP, the cGAS-generated second messenger, in HCC827OR versus parental cells assayed by LC-MS/MS analysis. (F) Western blot analysis of cGAS and STING proteins in HCC827OR versus parental cells. (G) Upregulation of p-STING in resistant cells, assayed by western blot analysis. (H) Effect of STING siRNA treatment (20 nM, 48 h) on cell growth in H1975OR versus parental cells. The cell growth was monitored and quantified using IncuCyte ZOOM system every 4 h. Mock siRNA at 20 nM as control. (I) Effect of STING knockdown (20 nM siRNA, 72 h) on cell viability in TKI-resistant cells versus parental cells. STING siRNA versus mock siRNA: ns, not significant; *, p < 0.05; **, p < 0.01; ***, p < 0.001; ****, p < 0.0001. (J) Effect of STING knockdown (10 nM siRNA, 48 h) on colony formation for 7 days in HCC827 resistant (ER, GR, OR) cells versus parental cells. Cells at a density of 1,000/well in 6-well plate were seeded at the beginning of the assay. Resistant versus parental: #, p < 0.05; ##, p < 0.01; ### p < 0.001; ####, p < 0.0001. STING siRNA versus mock siRNA: *, p < 0.05; **, p < 0.01; ***, p < 0.001; ****, p < 0.0001. (K) Effect of STING siRNA treatment on cell migration in HCC827 resistant (ER, GR, OR) cells versus parental cells. After 20 nM STING or mock siRNA treatment for 48 h, the cells were trypsinized, resuspended, and adjusted to 50,000/well in transwell for incubation of 24 h for cell migration. Resistant versus parental: ##, p < 0.01. STING siRNA versus mock siRNA: *, p < 0.05; **, p < 0.01. Representative images are demonstrated in Supplementary Fig. S2H. (L) Western blot analysis of epithelial marker E-cadherin (E-cad) and mesenchymal marker vimentin (vim). C, control, M, mock. (M and N) Effect of STING knockdown (M, 10 nM siRNA for 72 h) or inhibition (N, 10 μM H151 for 72 h) on TKI sensitivity in TKI-resistant cells. Combination index was calculated according to Chou Talalay method as described in Methods. (O and P) Effect of STING knockdown (O, 20 nM siRNA for 72 h) or inhibition (P, 10 μM H151 for 72 h) on cisplatin sensitivity in H460 human large cell lung carcinoma cells and C33A human cervical cancer cells.

Consistently, the resistant cells showed an intracellular accumulation of cytosolic DNA (Fig. 1C and D; Supplementary Fig. S1D), the endogenous cGAS agonist, and cGAMP (Fig. 1E), the cGAS-generated second messenger. Accordingly, the active form of cGAS and STING (cGAS dimers and STING dimers, respectively) (Fig. 1F) and the activated STING, phosphorylated STING (p-STING) (Fig. 1G) were accordingly upregulated in resistant cells. The activation of cGAS-STING was reconfirmed by RT-qPCR analysis showing the upregulation of signaling downstream gene expression (Supplemental Fig. S1E and S1F, red bar vs. grey bar), which depended on STING (Supplemental Fig. S1F).

### Addiction of resistant cells to STING pathway

These STING-activated, drug-resistant cells were more sensitive to STING knockdown or inhibition compared with their parental cells, assayed by the cell growth monitoring (Fig. 1H; Supplemental Fig. S2A and S2B) and viability (Fig. 1I; Supplemental Fig. S2C) analyses. Meanwhile, resistant cells demonstrated enhanced colony formation ability (Fig. 1J; Supplemental Fig. S2D) and epithelial-mesenchymal transition (EMT) properties including EMT gene enrichment (Supplemental Fig. S2E), epithelial marker and mesenchymal markers change (Supplemental Fig. S2F), morphologic transition (Supplemental Fig. S2G, OR vs. P), and migration ability (Fig. 1K; Supplemental Fig. S2H and S2I). These enhanced drug resistance properties were abrogated by STING knockdown (Fig. 1K, L; Supplemental Fig. S2G-S2I).

### Resistance to targeted and conventional chemotherapy drugs depends on STING activation

Knockdown of STING, either transiently by small interfering RNA (siRNA; Supplementary Fig. S3A) or constantly by short hairpin RNA (shRNA; Supplementary Fig. S3B and S3C), resensitized EGFR TKI-resistant cells to the corresponding TKI osimertinib (Fig. 1M; Supplementary Fig. S3D and S3E), erlotinib (Supplementary Fig. S3F), and gefitinib (Supplementary Fig. S3G). The resensitization was recapitulated by STING selective inhibitor H151 (Fig. 1N). Moreover, STING knockdown (Fig. 1O; Supplementary Fig. S3H-S3J) or inhibition (Fig. 1P) sensitized H460 large cell lung carcinoma cells, MCF-7 breast cancer cells, SW620 colon cancer cells, C33A cervical cancer cells, and the KRAS-G12S mutated, non-targetable A549 lung cancer cells to chemotherapy drug cisplatin, indicating that cancer cells refractory to conventional chemotherapy drugs depend on STING activation.

### Targeted or conventional chemotherapy drug treatment triggers cGAS-STING pathway activation

We then investigated how the cGAS-STING signaling was activated. Treatment with the targeted therapy drug osimertinib induced an intracellular accumulation of cytosolic dsDNA in sensitive cells (Fig. 2A), coincided with an induction of DNA damage as shown by an upregulation of the DNA double-strand break marker γ-H2AX (Fig. 2B). Consistently, accumulation of intracellular cGAMP (Fig. 2C) and upregulation of cGAS dimers and STING dimers (Fig. 2D) were demonstrated in these cells treated with osimertinib.

**Figure 2.**
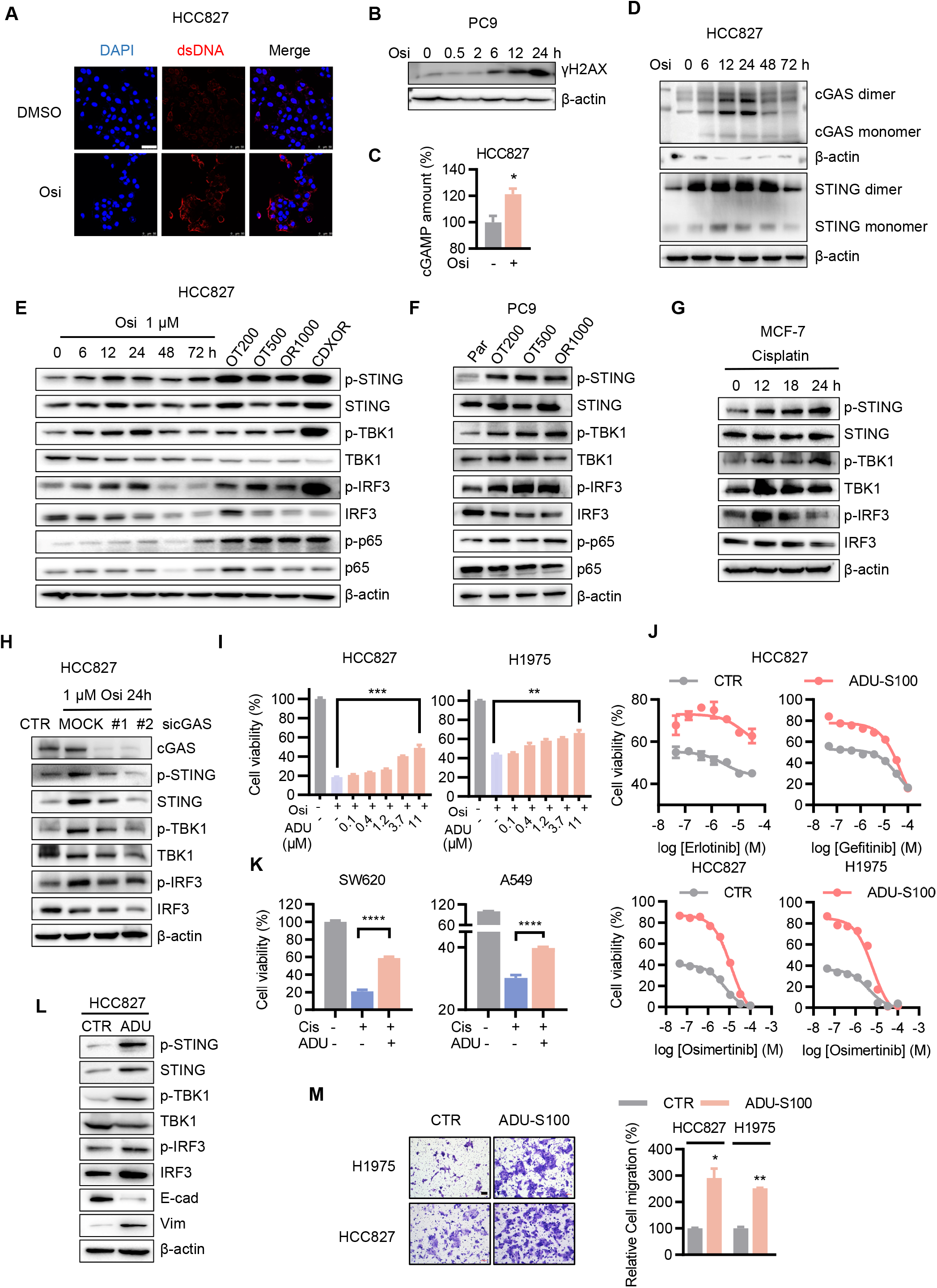
Drug treatment-triggered activation of cGAS-STING signaling is sufficient for resistance induction. (A) Immunofluorescence staining analysis of dsDNA accumulation in the cytoplasm of HCC827 cells exposed to TKI acute treatment (1 μM osimertinib for 2 h). Scale bar: 50 μm. (B) Western blot analysis of DNA damage marker γH2AX in PC9 cells exposed to 1 μM osimertinib as indicated period. (C) LC-MS/MS analysis of intracellular cGAMP in HCC827 cells exposed to 1 μM osimertinib for 72 h. (D) Western blot analysis of cGAS dimer and STING dimer in HCC827 cells exposed to 1 μM osimertinib as indicated period. (E) Western blot analysis of cGAS-STING signaling activation markers p-STING, p-TBK1, p-IRF3, and p-p65 in the process of resistance emergence and maintenance (sensitive, tolerant, and resistant/relapsed phases) in in vitro and in vivo models. HCC827 cells in the sensitive phase were exposed to 1 μM osimertinib as indicated period. OT200 and OT500 are HCC827-derived cell lines that are tolerant to 200 and 500 nM osimertinib, respectively. OR1000 is the HCC827-derived cell line that is resistant to 1,000 nM osimertinib. CDXOR is the HCC827CDX tumor in BALB/c nu/nu athymic mice relapsed to osimertinib (2 mg/kg, p.o., q.d.) treatment. (F) Western blot analysis of cGAS-STING signaling activation markers p-STING, p-TBK1, p-IRF3, and p-p65 in PC9 tolerant and resistant model cell lines to osimertinib. OT200 and OT500 are PC9-derived cell lines that are tolerant to 200 and 500 nM osimertinib, respectively. OR1000 is the PC9-derived cell line that is resistant to 1000 nM osimertinib. (G) Western blot analysis of cGAS-STING signaling activation markers in MCF-7 cells exposed to 100 μM cisplatin as indicated periods. (H) Effect of cGAS knockdown on osimertinib-induced STING signaling activation. After incubated with 20 nM cGAS or mock siRNA for 24 h, the cells were further exposed to 1 μM osimertinib for 24 h. (I) Concentration-dependent effect of STING agonist ADU-S100 on osimertinib sensitivity in HCC827 and H1975 cells. After incubated with indicated concentrations of ADU-S100 for 24 h. The cells were exposed to 1 μM osimertinib for 48 h. (J and K) Effect of ADU-S100 on TKIs (J) and cisplatin (K) sensitivity. After incubated with 10 μM ADU-S100 for 24 h, the cells were exposed further to indicated concentrations of individual TKIs or 33 μM cisplatin for 48 h. (L) Effect of ADU-S100 (10 μM for 24 h) on STING signaling and EMT markers in HCC827 cells. (M) Effect of ADU-S100 on H1975 and HCC827 cell migration. After 10 μM ADU-S100 treatment for 24 h, the cells were trypsinized, resuspended, and adjusted to 50,000/well in transwell for incubation of 24 h for cell migration. *, p < 0.05; **, p < 0.01.

Consequently, cGAS-STING signaling was triggered, evidenced by a time-dependent increase of activated form of pathway regulators and effectors p-STING, p-TBK1, p-IRF3, and p-p65 after osimertinib treatment (Fig. 2E). The activation sustained throughout the whole process of resistance emergence and maintenance in in vitro and in vivo models (Fig. 2E and F). This drug stress-induced STING downstream signaling activation was not confined to targeted therapies, as evidenced by an activation of STING signaling in breast cancer MCF-7 cells underwent cisplatin treatment (Fig. 2G), extending to chemotherapy situations. The activation was abrogated by cGAS suppression (Fig. 2H), indicating its dependence on cGAS.

### STING activation is sufficient for resistance induction

Exogenous supplement of cGAMP, the endogenous STING agonist, induced the otherwise sensitive cells resistant to TKIs (Supplementary Fig. S3K). This resistance-induction effect of STING activation was recapitulated by more cell membrane-permeable STING agonist ADU-S100 (Fig. 2I) treatment in targeted therapy (Fig. 2J; Supplementary Fig. S3L) and in chemotherapy situations (Fig. 2K). Moreover, ADU-100 administration was sufficient to induce EMT, a key property for resistance, as demonstrated by marker change (E-cad downregulation and Vim upregulation) (Fig. 2L) and migration enhancement (Fig. 2M).

### STING confers resistance via activation of TBK1-IRF3/p65 NF-κB signaling

Transcriptomic analysis showed inflammatory properties in EGFR TKI-resistant tumors (Fig. 3A; Supplementary Fig. S4A) and cells (Supplementary Fig. S4B). Consistently, analysis of public clinical data recorded in The Cancer Genome Atlas (TCGA) database showed that in targeted and chemotherapy drug-treated lung cancer patients, inflammatory response properties were enriched in tumors with EMT features (Supplementary Fig. S4C), a well-known drug resistance mechanism (12). Western blot analysis demonstrated in resistant cells an upregulation of STING downstream inflammatory signaling factors p-STING, p-IRF3, and p-p65 (Fig. 2E, 2F, and 3B) which depended on STING (Fig. 3C). IRF3 and p65 were important for resistant cell survival (Fig. 3D), growth (Supplementary Fig. S4D), EMT/migration (Fig. 3E and F; Supplementary Fig. S4E and S4F), and insensitivity to TKIs (Fig. 3G and H; Supplementary Fig. S4G-S4L). STING activation depended on IRF3 and p65 for drug resistance induction, as shown by the fact that the STING activation-induced osimertinib resistance was completely abrogated by the combination of IRF3 knockdown and p65 knockdown (Fig. 3I).

**Figure 3.**
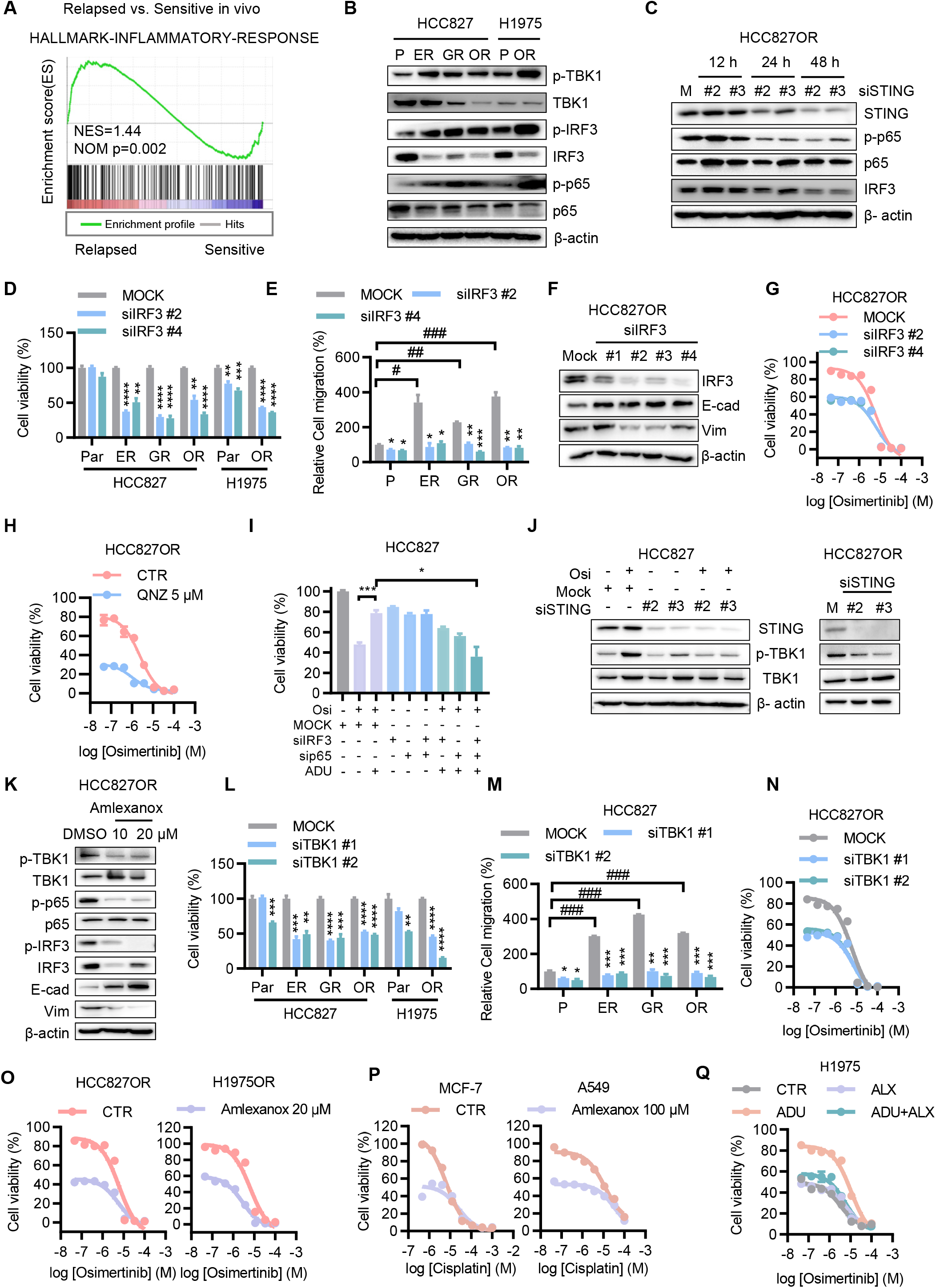
STING confers resistance via activation of TBK1-IRF3/NF-κB signaling. (A) GSEA of RNA-seq data indicates an inflammatory hallmark in erlotinib treatment-relapsed tumors (n=5) compared with the sensitive tumors (n=6). (B) Western blot analysis of STING downstream inflammatory signaling markers p-TBK1, p-IRF3, and p-p65 in resistant (ER, GR, OR) versus parental (P) cells. (C) Effect of STING knockdown (20 nM siRNA) on STING downstream inflammatory signaling in HCC827OR cells. (D) Selective effect of IRF suppression (20 nM siRNA for 72 h) on resistant cell viability. (E) Selective effect of IRF suppression (20 nM siRNA for 48 h) on enhanced migration in HCC827ER, GR, and OR resistant cells (representative images in Supplementary Fig. S4E). After IRF siRNA or mock siRNA treatment for 48 h, the cells were trypsinized, resuspended, and adjusted to 50,000/well in transwell for incubation of 24 h for cell migration. Resistant versus parental: #, p < 0.05; ##, p < 0.01; ###, p < 0.001. siIRF3 versus mock: *, p < 0.05; **, p < 0.01; ***, p < 0.001. (F) Effect of IRF3 suppression (20 nM siRNA for 48 h) on resistant cell EMT markers. (G and H) Effect of suppression of IRF3 (G, 10 nM siRNA for 72 h) or p65 (H, 5 μM QNZ for 72 h) on osimertinib concentration-cell viability inhibition response curves in HCC827OR cells. (I) Effect of a combination of IRF knockdown and p65 knockdown on the STING activation-induced osimertinib resistance. The cells were incubated with 10 μM ADU-S100 for 24 h, and exposed further to 20 nM IRF siRNA, 20 nM p65 siRNA, 1 μM osimertinib alone, or in combination as indicated for 48 h. (J) Effect of STING knockdown on osimertinib acute treatment-induced TBK1 activation in HCC827 parental cells (left panel) and on the constitutively activated TBK1 in HCC827OR cells (right panel). Parental cells were treated with 20 nM STING siRNA for 48 h and then exposed to 1 μM osimertinib for 2 h; OR cells were treated with 20 nM STING siRNA for 72 h. (K) Effect of TBK1 suppression (amlexanox for 48 h) on cGAS-STING signaling and EMT markers in HCC827OR cells. (L and M) Effect of TBK1 suppression (20 nM siRNA) on resistant cell viability (L) and enhanced migration (M, representative images in Supplementary Fig. S5C). siRNAs were used for 72 h in viability assay and for 48 h in the migration assay. (N and O) Effect of TBK1 knockdown (N, 10 nM siRNA, 72 h) or inhibition (O, amlexanox, 72 h) on osimertinib sensitivity in HCC827OR and H1975OR resistant cells. (P) Effect of TBK1 suppression (amlexanox, 72 h) on cisplatin sensitivity in MCF-7 and A549 cells. (Q) Effect of TBK1 suppression (20 μM amlexanox, ALX) on STING activation (10 μM ADU-S100)-induced osimertinib resistance. After incubated with ADU-S100 for 24 h, the cells were further exposed to osimertinib with or without ALX for 48 h.

STING instigated IRF3 and p65 signaling via TBK1. Targeted and chemotherapy drugs induced an activation of TBK1 in sensitive cells (Fig. 2E and G) and the activation was maintained in resistance phase (Fig. 2E, 2F, and 3B). Both the TKI-induced activation of TBK1 in sensitive cells (Fig. 3J, left panel) and the constitutive activation of TBK1 in resistant cells (Fig. 3J, right panel) were abrogated by STING knockdown. This TBK1 activation controlled downstream IRF3 and p65, evidenced by a nullification of upregulated p-IRF3 and p-65 in resistant cells by TBK1 selective inhibitor amlexanox (Fig. 3K).

Then we examined if TBK1 mediated the STING-induced drug resistance and cell EMT/migration enhancement. Genetic knockdown (Supplementary Fig. S5A) or pharmacological inhibition of TBK1 selectively suppressed the resistant cell viability (Fig. 3L; Supplementary Fig. S5B) and EMT/migration properties (Fig. 3M; Supplementary Fig. S5C and 5D) and resensitized these cells to targeted (Fig. 3N and O; Supplementary Fig. S5E) and chemotherapy (Fig.3P; Supplementary Fig. S5F) drugs. Additionally, TBK1 suppression abrogated the STING activation-induced osimertinib resistance (Fig. 3Q; Supplementary Fig. S5G) and migration enhancement (Supplementary Fig. S5H).

### Drug resistance acquisition and maintenance in vivo depend on STING activation

STING signaling was triggered and activated by drug treatment and the activation was maintained throughout the whole process of resistance emergence and maintenance (Fig. 2E-G) and this activation was required and sufficient for drug resistance in vitro. We next examined whether the role of STING signaling in drug resistance was recapitulated in vivo in immunodeficient nude mice. HCC827 cell-derived xenograft (CDX) tumors initially responded well to osimertinib with growth delay and volume decrease after osimertinib administration but resistance was acquired after continuous drug treatment as shown by a tumor burden relapse (Fig. 4A). Blockade of STING signaling with amlexanox not only prevented the acquisition of resistance but also resensitized the resistance-acquired tumors to osimertinib (Fig. 4 A, B, and C), indicating that both the resistance acquisition and maintenance depend on STING signaling in vivo. STING signaling blockade with amlexanox did not cause mouse body weight change (Supplementary Fig. S5I).

**Figure 4.**
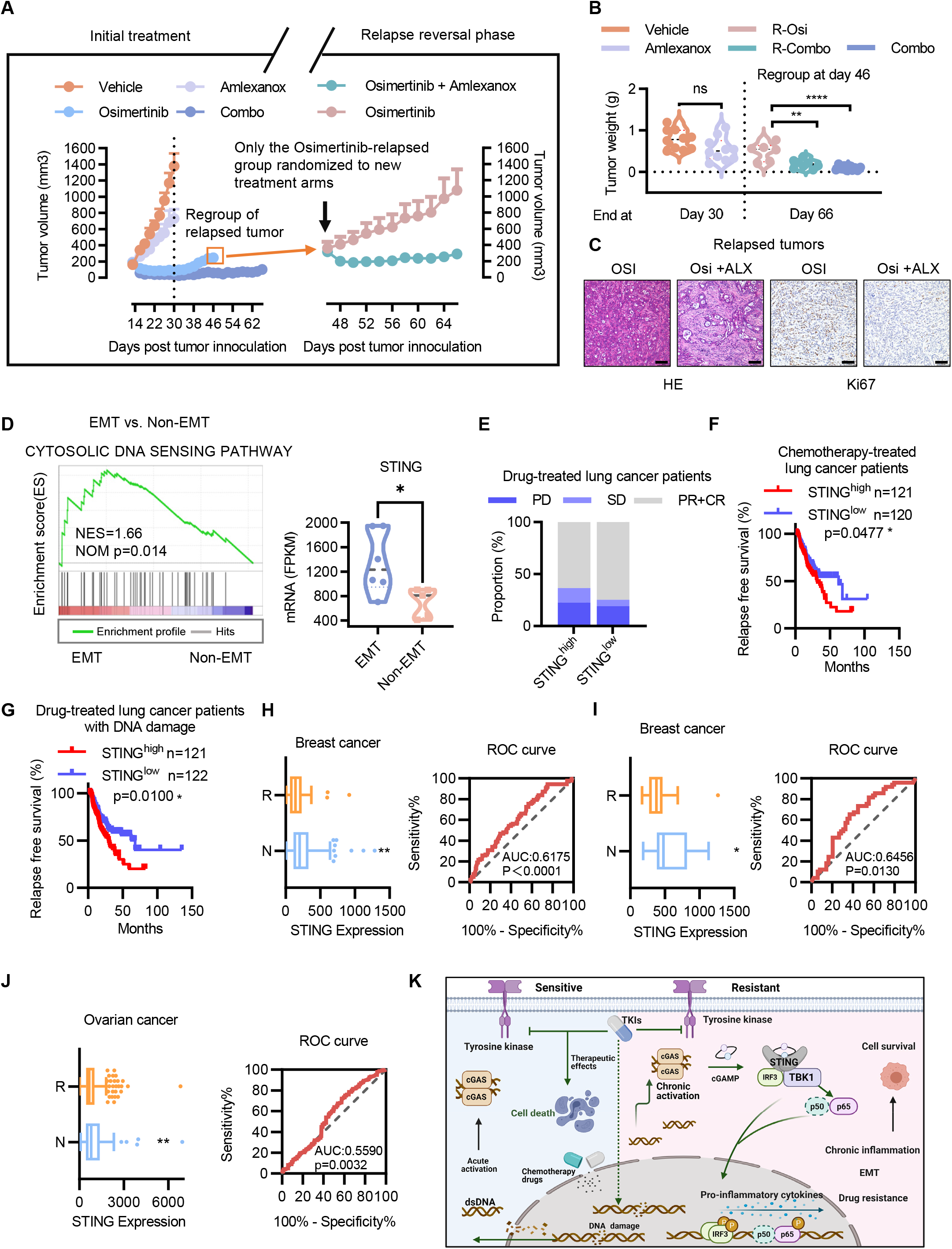
Drug resistance acquisition and maintenance in vivo depend on STING signaling. (A and B) Effect of STING signaling blockade with amlexanox on prevention and reversal of osimertinib resistance in HCC827CDX tumors. Shown are tumor growth over time (A) and tumor weights at day 30 and day 66 (B). R-Osi represents continued osimertinib single agent treatment for relapsed tumors; R-Combo represents osimertinib + amlexanox combination treatment for relapsed tumors; Combo represents the osimertinib + amlexanox combination treatment throughout the whole experiment). Flank subcutaneously implanted, HCC827 cell-derived xenograft tumors (2 tumors per mouse), when reached ~ 200 mm^3^, were randomized into groups, at day 13: Vehicle control (n=5), Amlexanox (n=5), Osimertinib (n=10), and Combo (combination of amlexanox and osimertinib) (n=5). At day 46 post tumor inoculation, 7 mice in osimertinib group relapsed and were randomly regrouped into Osimertinib (continuation) arm (n=3) and Amlexanox + Osimertinib arm (n=4). STING signaling pathway inhibitor amlexanox and EGFR TKI osimertinib were administered (p.o., q.d.) as 50 mg/kg and 2 mg/kg, respectively. (C) Representative images of hematoxylin and eosin (H&E) staining for pathological analysis and immunohistochemistry (IHC) staining of the cell proliferation marker Ki-67 for in situ analysis of cell proliferation. Scale bar: 100 μm. (D) GSEA of RNA-seq data indicates an increased activation of cytosolic DNA sensing pathway in tumors with EMT versus tumors with non-EMT signature in lung cancer patients that have undergone targeted or chemotherapy drug treatment (Six patients have tumors with prototypical EMT signature [Vimentin expression abundance ranks top 10% and E-cadherin ranks bottom 10% in total tumors] and 5 with prototypical non-EMT signature [Vimentin ranks bottom 10% and E-cadherin ranks top 10%]. Two hundred eighty-eight patients who had drug treatment information in TCGA data were analyzed. Vimentin and E-cadherin are mesenchymal and epithelial maker, respectively. (E) Response profiles of lung cancer patients underwent EGFR TKI (n=24) or chemotherapy (n=206) treatment. CR, complete response. PR, partial response. SD, stable disease. PD, progressive disease. (F and G) Kaplan-Meier analysis of relapsed shows that lower free survival (RFS) probability was correlated with higher tumor STING expression in lung cancer patients underwent chemotherapy (F) and in lung cancer patients who underwent targeted and chemotherapy drug treatment as well as got DNA damage in their tumors (G). In E, F, G, data were retrieved from TCGA data set (http://cancergenome.nih.gov/). The patients were stratified according to high versus low expression (cutoff: median) of STING mRNA abundance within their tumors. (H-J) Box-plots and ROC (receiver operating characteristic) curves of predictive effect of STING on chemotherapy response defined as RFS in ovarian cancer (H), pathological response (I) and RFS (J) in breast cancer. The analyses were performed according to the ROC Plotter tool (http://www.rocplot.org/). (K) Schematic demonstration of the drug resistance conferred by cancer cell-autonomous cGAS-STING response to drug treatment stress. Created with BioRender.com.

Consistently, analysis of public clinical data recorded in TCGA database showed that in targeted and chemotherapy drug-treated lung cancer patients, cytosolic DNA sensing pathway was enriched in tumors with EMT features (Fig. 4D). Meanwhile, the patients whose tumors expressed higher abundance of STING demonstrated a lower drug response (Fig. 4E) and relapse-free survival (RFS) probability (Fig. 4F). In patients who underwent drug treatment and got DNA damage in tumors, higher expression of STING correlated with lower RFS probability (Fig. 4G). Additionally, lung cancer patients with enhanced cGAS-STING signaling signature demonstrated lower overall survival probability (Supplementary Fig. S5J).

The negative effect of STING on drug response was recapitulated in breast (Fig. 4H and I) and ovarian ((Fig. 4J) cancers, where higher expression of STING predicted inferior response to chemotherapies.

## Discussion

Recent progress has uncovered the multifaceted roles of STING signaling in cancer biology, not only triggering anticancer immune response around the tumor but also inducing tumorigenesis and metastasis (1,2).

In this study, we find a new face of cGAS-STING signaling whose activation in a cancer-cell-autonomous response manner confers cancer drug resistance, highlighting a potent ability of cancer cells hijacking the evolutionarily conserved inflammatory signaling to copy with drug stress.

Anticancer drugs that act primarily through DNA-damaging mechanism, such as etoposide, cisplatin, and PARP inhibitors, have been reported to trigger cytosolic DNA accumulation and cGAS-STING signaling activation (3,13). Besides chemotherapy drugs, the molecularly targeted drug EGFR TKIs, acting primarily not as traditional chemotherapy cytotoxic agents, have been demonstrated in this study, also induced cancer cell cytosolic DNA accumulation and triggered the cell-intrinsic cGAS-cGAMP-STING activation and downstream signaling transduction.

STING activation was sufficient and necessary for acquisition and maintenance of the cancer drug resistance. Activation of STING induced drug resistance acquirement, which was abrogated by STING antagonism in vitro; blockade of STING signaling prevented the acquired resistance and the subsequent tumor relapse to therapy in vivo, indicating the importance of STING activation on drug resistance emergence. On the other hand, STING activation also plays important roles in maintenance of the resistance. First, resistant cells exhibited a constitutive hyperactivation of STING signaling which is pivotal for survival and withstanding drug insult demonstrated by the fact that genetic or pharmacological blockade of STING signaling selectively suppressed the resistant cells and resensitized the cells to drug treatment. Second, in vivo, drug-resistant and relapse were overcome by blockade of STING signaling.

STING conferred resistance through its TBK1-IRF3/NF-κB downstream signaling. TBK1 stands at the hub of STING signaling and inflammatory responses and is essential to cancer cell survival and growth (14,15). Moreover, recent discoveries have identified TBK1 as a key point and a therapeutic target for cancer drug resistance (16,17) and the activation of TBK1-IRF3 signaling has been reported to protect cells from drug-induced death (18). We found that the STING-conferred drug resistance depended on its downstream p65-relied canonical NF-κB signaling. Effect of canonical NF-κB signaling on drug resistance induction has been reported (19). STING-induced metastasis, in contrast, was mediated by noncanonical NF-κB signaling (4). It is intriguing that cancer metastasis and drug resistance are mediated by divergent pathways of STING downstream signaling, deserving further investigation.

Taken together, targeted or conventional chemotherapy drug-induced activation of cGAS-STING in cancer cells confers these cells an ability to withstand the drug challenge in a cancer cell-adapted and autonomous response manner. DNA directly or indirectly damaged by drug pressure leaks into the cytosol. The accumulated cytosolic DNA is intrinsically sensed and responded by cGAS-cGAMP-STING signaling pathway. Cancer cells can take advantage of this pathway via the downstream TBK1-IRF3/NF-κB signaling to counteract drug stress and acquire resistance to these drugs (Fig. 4K).

This finding has important implications for a better understanding of cGAS-STING functions and cancer therapies. First, a new face of STING signaling and a novel skill of cancer cells autonomously adapted for counteracting drug stress are uncovered and formally proposed. Second, the cancer-cell-autonomous effects of the cell-intrinsic cGAS-STING activation, other than the modulation of microenvironmental immune cells, are worthy of attention. Third, there are numerous STING agonists being clinically tested, either as a single agent or in combination with targeted or chemotherapy drugs (20), our discovery of STING activation as a drug resistance-inducing mechanism implies a possible explanation, at least partly, for their limited efficacy and warrant a caution in designing these kinds of combination strategies.

## Methods

### Cells and Cell Culture

Human lung adenocarcinoma cell lines that harbor EGFR-activating mutations, HCC827 and H1975, were obtained from American Type Culture Collection (ATCC), PC9 from Dr. G.L. Zhuang (China State Key Laboratory of Oncogenes and Related Genes). Human large cell lung cancer cell line H460, human lung adenocarcinoma cell line A549, human cervical cancer cell line C33A, human breast cancer cell line MCF7, and human colorectal cancer cell line SW620 were obtained from Dr. Q. Lu (Department of Pharmacology and Chemical Biology, Shanghai Jiao Tong University School of Medicine). The cells were authenticated by short tandem repeat (STR) analysis. Cell lines which are resistant to erlotinib (HCC827ER), gefitinib (HCC827GR), and osimertinib (HCC827OR and H1975OR1–5) were established, maintained, and authenticated as schematized (Supplementary Fig. 1A) and previously described (9–11). The cells were cultured in RPMI 1640 medium (Gibco, USA) containing 10% FBS, 1% GlutaMAX, and 1% penicillin-streptomycin at 37 °C with 5% CO2.

### Reagents and antibodies

Erlotinib and gefitinib were purchased from LC Laboratories. Osimertinib, amlexanox, QNZ (EVP4593) were purchased from Selleck Chemicals. ADU-S100 were purchased from MCE. Dimethyl sulfoxide (DMSO) was obtained from Sigma. Antibodies for dsDNA and p-IRF3 were purchased from Abcam; for E-cadherin, vimentin, β-actin, STING, p-STING, TBK1, p-TBK1, p65, p-p65, IRF3 and γH2AX from Cell Signaling Technology; for cGAS from Sigma.

### Cell Growth and Viability Assay

Cells were seeded in 96-well plates at a density of 4,000 cells/well. After the cells adhering to the well, corresponding treatment agents were added to the medium. Cell growth was monitored using the IncuCyte ZOOM live cell analysis system (Essen Bioscience). Cell viability was determined using the Cell Counting Kit-8 (CCK-8; Dojindo) assay according to the manufacturer’s instructions.

### Colony Formation Assay

Cells were seeded in 6-well plates (1,000 cells/well) for treatments and cultured in complete medium for about 1 week. Fixed with 4% paraformaldehyde for 15 minutes, the cells were stained with 0.01% crystal violet for counting. Individual colonies (>50 cells) were counted.

### Transwell Migration Assay

Migration assays were performed by using Transwell chambers (Corning Costar). After the corresponding treatments, the cells (50,000 per chamber) were plated into the upper chambers containing serum-free medium whereas the lower chambers were filled with medium containing 10% FBS. After 24 hours, the cells attached to the underside of the filter were stained with 0.1% crystal violet solution and imaged (Nikon). Then, the stained cells were dissolved with 10% acetic acid (100 μL/chamber), and the optical density (OD) was detected at 600 nm using a microplate reader to quantify the cell migration ability.

### Western Blot and Immunofluorescence Analysis

Western blot and immunofluorescence analyses were performed according to standard protocols. First antibodies are described in Methods or indicated in corresponding figure legends.

### Gene knockdown by RNA interference

For transient knockdown, cells were transfected with small interfering RNAs (siRNAs) using Lipofectamine 3000 reagent (Invitrogen, USA) according to the manufacturer’s instructions. Target sequences were used as shown in Supplementary Table S1. For constant knockdown, endogenous STING was silenced by using short hairpin RNA (shRNA), with scrambled shRNAs as controls. Shortly, pGIPZ-shSTING expression clones were packaged with pMD2.G and pSPAX2. For transfection, cells were seeded in 6-well plates at density of 200,000/well. Twenty-four hours later, the lentiviral particles were diluted with serum-free medium containing 6 μg/mL polybrene. The sense sequences of shRNAs were shown in Supplementary Table S1.

### Liquid Chromatography and Tandem Mass Spectrometry Analysis of cGAMP

Cell samples were analyzed on an Exion^®^ LC-system coupled with a TripleQUAD^®^ 6500 plus system (Sciex, Concord, ON, USA). A volume of 5 μL was injected onto a BEH HILIC UPLC column (1.7 μm, 100 × 3 mm; Waters). The column oven was 35 °C. The mobile phase was set as follows: (A) 10 mM ammonium fluoride and 0.1% ammonia in water, (B) pure acetonitrile. The initial condition was 45% A, maintained for 1 min. The mobile phase was ramped to 72% A from 1 min to 7 min, then maintained at 45% A from 7.01 min to 10 min. The flow rate was set to 0.4 ml/min. The mass spectrometer was operated in Electron Spray Ionization (ESI) negative mode with the source temperature 550 °C. Declustering and collision-induced dissociation were achieved with nitrogen gas. The multiple reaction monitoring (MRM) transition, declustering potential (DP) and collision energy for cGAMP measurement were as follows: m/z 673.1 [M-H]- → 344.1, −200 V, −50 eV The analytic data were processed using AnalystTM 1.7 software and MultiQuantTM 3.0 software (Sciex, Concord, ON, USA).

### Animal Study and In Vivo Xenografts Assay

Animal work was evaluated and approved by the Institutional Animal Care and Use Committee (IACUC) of Shanghai Jiao Tong University School of Medicine. In cell-derived xenograft (CDX) studies, one hundred microliters of phosphate buffered saline (PBS) containing 5 × 10^6^ cancer cells were subcutaneously injected into the left and right flanks of 4-week-old BALB/c nu/nu athymic mice. Subcutaneous local tumors were measured on length (L) and width (L) by a Vernier caliper every 2-3 days and the volume of tumors were calculated with the formula: V = LW^2^/2. When tumor volume reached ~ 200 mm^3^, mice were randomized into different treatment groups as described in figure legends. Mice were euthanized when the animal experiments reached the end or the tumor volume of the vehicle control group reached 1,500 mm^3^.

### Combination Effect Analysis

Treatment combinations will lead to synergy, antagonism, or additive effect. A calculation of Combination Index (CI) based on Chou Talalay method (21) (Chou06PharmacoRev) was used to evaluated the combination effects. CompuSyn software (CompuSyn Inc. USA) was utilized to analyze the data.

### Statistical analysis

Quantitative data are presented as mean ± SEM of at least three independent experiments. Differences were assessed using the two-tailed Student’s t test or ANOVA with Bonferroni posttest unless otherwise indicated. P < 0.05 was considered statistically significant. *, P < 0.05; **, P < 0.01. ***, P < 0.001; ****, P < 0.0001 unless otherwise indicated.

## Acknowledgements

This work was supported by grants from the National Natural Science Foundation of China (No. 81872882, 81573018, 82003772) and Shanghai Municipal Science Foundation (No. 21ZR1436700).

## Author contributions

Q.-M.L. and H.-M.L. established in vitro and in vivo experiment conditions. Q.-M.L. performed analyses for cell growth, viability, western blot, and in vivo response under drug treatment. Q.-M.L. performed analyses for TCGA, RNA-seq, and omics data mining. S.-Y.W. and K.-R.Z. cultured cells, performed analyses for cell colony formation and apoptosis. Y.S. provided study materials and contributed to manuscript writing and revision. L.-Q.M., S.-Y.W., and H.-M.L. designed the study, analyzed the data, and contributed to manuscript writing and revision. L.-M.L and H.-Z.C interpreted data, acquired financial support, and contributed to manuscript writing and review. H.-Z.C contributed to oversight and leadership responsibility for the research and approved the manuscript. L.Z designed study conception and experiments, assembled and interpreted data, wrote the manuscript, and approved the manuscript.

