## Supplementary Information for "Cancer cell-autonomous cGAS-STING response confers drug resistance"

**Figure S1. Hyperactivation of cGAS-STING signaling in drug-resistant cancer cells**

(A) Schematic overview of the establishment of in vitro (top panel) and in vivo (bottom panel) EGFR TKI-resistant models.

(B) Gene set enrichment analysis (GSEA) of RNA-seq data indicates a hyperactivation of cytosolic DNA sensing pathway in EGFR TKI-resistant (HCC827GR, PC9ER, H1975OR) and cisplatin-resistant (A549DR) cells compared with their corresponding isogenic parental cells. GR, ER, OR, and DR represent gefitinib, erlotinib, osimertinib, and DDP (cis-diamminedichloroplatinum, cisplatin) resistance, respectively.

(C) Heatmap shows expression difference of cGAS-STING signaling downstream genes between H1975OR1-5 and parental (Par) cells assayed by RNA-seq analysis. H1975-OR1, -OR2, -OR3, -OR4, and -OR5 cells were H1975 resistant cells that derived from those grew in osimertinib with the final concentrations of 1, 2, 3, 4, and 5 μM, respectively.

(D) Double-strand DNA (dsDNA) accumulation in the cytoplasm of HCC827GR and PC9OR resistant versus corresponding parental (P) cells assayed by anti-dsDNA antibody-based immunofluorescence staining analysis. Scale bar: 50 μm.

(E and F) RT-qPCR analysis of STING downstream gene expression in H1975OR versus H1975 cells (E) and HCC827OREV versus HCC827EV versus HCC827OR-shSTING cells (F). HCC827OR-shSTING cells are HCC827OR cells where STING has been constantly knocked down by short hairpin RNA (shRNA). EV, empty vector control.

**Figure S2. Addiction of resistant cells to STING pathway**

(A-B) Selective effect of STING siRNA treatment (20 nM, 48 h) on cell growth in HCC827OR (A) and H1975OR (B, Scale bar: 600 μm) versus parental cells. The cell growth was monitored and quantified using IncuCyte ZOOM system every 4 h.

(C) Effect of STING inhibition (H151 for 72 h) on cell viability in TKI-resistant cells versus parental cells.

(D) Effect of STING knockdown (10 nM siRNA, 48 h) on colony formation for 7 days in HCC827 resistant (ER, GR, OR) cells versus parental cells. Cells at a density of 1,000/well in 6-well plate were seeded at the beginning of the assay.

(E) GSEA of RNA-seq data indicates an EMT hallmark in HCC827GR compared with parental cells.

(F) EMT in resistant cells (HCC827ER, GR, OR vs parental; H1975OR vs parental) assayed by western blot analysis of EMT markers.

(G) EMT morphology properties (top panel) assayed by bright-field analysis (Scale bar: 300 μm) and the effect of STING knockdown with shRNA on EMT morphology (bottom panel, Scale bar:100 μm)

(H-I) Effect of STING siRNA treatment on cell migration in HCC827ER, GR, OR (H) and H1975OR (I) cells versus parental cells. After 20 nM STING or mock siRNA treatment for 48 h, the cells were trypsinized, resuspended, and adjusted to 50,000/well in transwell for incubation of 24 h for cell migration. Resistant versus parental: ##, p <

0.01. STING siRNA versus mock siRNA: *, $p < 0.05$; **, $p < 0.01$; ****, $p < 0.0001$. ns, not statistically significant.

**Figure S3. STING activation is required and sufficient for resistance**

(A-C) Western blot (A and B) and RT-qPCR (C) analyses of STING expression after transient (A) or constant (B and C) knockdown of STING by small interfering RNA (siRNA) and short hairpin RNA (shRNA), respectively.

(D and E) Effect of constant knockdown of STING on HCC827OR cell sensitivity to osimertinib treatment (72 h) assayed by cell viability (D) and growth (E) analyses.

(F and G) Effect of transient knockdown of STING (10 nM siRNA, 72 h) on HCC827ER (F) and HCC827GR (G) cell sensitivity to erlotinib and gefitinib, respectively.

(H-J) Effect of transient knockdown of STING (20 nM siRNA, 72 h) on A549 lung cancer (H), MCF-7 breast cancer (I), and SW620 colon cancer (J) cell sensitivity to cisplatin.

(K and L) Effect of STING agonists cGAMP (72 h) and ADU-100 on osimertinib sensitivity in H1975 and HCC827 cells (K) and on sensitivity to osimertinib, erlotinib, or gefitinib in PC9 cells (L). Two mg/l cGAMP (Sigma, USA) was transfected into cells using Lipofectamine 3000 reagent (Invitrogen, USA). After incubated with 10 μM ADU-S100 for 24 h, cells were further exposed to indicated concentrations of TKIs for 48 h.

**Figure S4. Resistant cells demonstrate inflammatory properties and dependence on IRF3 and p65.**

(A) Expression difference of cGAS-STING signaling downstream genes between erlotinib treatment-relapsed tumors (n=5) compared with the sensitive tumors (n=6) assayed by RNA-seq analysis.

(B and C) GSEA of RNA-seq data indicates inflammatory hallmarks in EGFR TKI-resistant (HCC827GR, PC9ER, H1975OR) versus their corresponding isogenic parental cells (B) and in tumors with EMT versus tumors with non-EMT signature in lung cancer patients that have undergone targeted or chemotherapy drug treatment (C). (Six patients have tumors with prototypical EMT signature [Vimentin expression abundance ranks top 10% and E-cadherin ranks bottom 10% in total tumors] and 5 with prototypical non-EMT signature [Vimentin ranks bottom 10% and E-cadherin ranks top 10%]. Two hundred eighty-eight patients who had drug treatment information in TCGA data were analyzed. Vimentin and E-cadherin are mesenchymal and epithelial maker, respectively.

(D) Selective effect of IRF3 knockdown (20 nM siRNA, 72 h) on HCC827OR versus HCC827 parental cell growth.

(E and F) Selective effect of IRF3 suppression (20 nM siRNA for 48 h) on enhanced migration in HCC827 (E) and H1975 (F) resistant cells. After IRF siRNA or mock siRNA treatment for 48 h, the cells were trypsinized, resuspended, and adjusted to 50,000/well in transwell for incubation of 24 h for cell migration. Resistant versus parental: ##, $p < 0.01$. siIRF3 versus mock: *, $p < 0.05$; **, $p < 0.01$.

(G-L) Resensitization effect of IRF3 knockdown (G-I, 10 nM siRNA, 72 h), p65 knockdown (J and K, 10 nM siRNA, 72 h), and p65 inhibition (L, 5 μM QNZ, 72 h) on TKI response in resistant cells.

**Figure S5. STING activation-induced resistance and migration depends on TBK1**

(A) Western blot analysis of TBK1 protein abundance in HCC827OR cells treated with TBK1- or mock- siRNAs (20 nM, 72 h).

(B) Selective effect of TBK1 inhibition (200 μM amlexanox, 72 h) on resistant cells versus corresponding parental cells.

(C and D) Selective effect of TBK1 suppression (20 nM siRNA, 48 h) on enhanced migration in HCC827 (C) and H1975 (D) resistant cells. After TBK1 siRNA or mock siRNA treatment for 48 h, the cells were trypsinized, resuspended, and adjusted to 50,000/well in transwell for incubation of 24 h for cell migration. Resistant versus parental: #, $p < 0.05$. siTBK1 versus mock: *, $p < 0.05$; **, $p < 0.01$.

(E and F) Sensitization effect of TBK1 knockdown (E, 10 nM siRNA, 72 h) or inhibition (F, 100 μM amlexanox, 72 h) on TKI response in HCC827ER and H1975OR cells and on cisplatin response in SW620 cells.

(G) Effect of TBK1 knockdown (20 nM siRNA) on STING activation (10 μM ADU-S100)-induced osimertinib resistance in HCC827 and H1975 cells. After incubated with ADU-S100 for 24 h, the cells were further exposed to osimertinib with TBK1 siRNA or mock siRNA for 48 h.

(H) Effect of TBK1 suppression (20 μM ALX) on STING activation (10 μM ADU-

S100)-induced migration in H1975 cells. After incubated with ADU-S100 or DMSO for 24 h, the cells were further exposed to 20 μM ALX or DMSO for 48 h. Then the cells were trypsinized, resuspended, and adjusted to 50,000/well in transwell for incubation of 24 h for cell migration.

(I) Analysis of body weight change.

(J) Kaplan-Meier analysis shows that lung cancer patients with enhanced cGAS-STING signaling signature (based on mRNAs abundance of cGAS, CCL2, CCL20, IL6, and TNF) in tumors demonstrated lower overall survival probability. Data were retrieved from TCGA LUAD and LUSC data sets (http://cancergenome.nih.gov/).

**Table S1. siRNA sequences for knockdown and primers for real-time qPCR.**

| | |
|---|---|
| Oligonucleotides | |
| siRNA | |
| Gene name | siRNA sequence |
| siSTING #1 | TGGTCATATTACATCGGAT |
| siSTING #2 | AGCATTACAACAACCTGCT |
| siSTING #3 | CGAACTTACAATCAGCATT |
| sicGAS #1 | TTCATATTCAATTTGCTTTGTC |
| sicGAS #2 | TTCTAAAAACTGACTCAGAGGA |
| siTBK1 #1 | AGAGATTTACTATCAGTTC |
| siTBK1 #2 | GCATTAAATGAATGCCTTT |
| siTBK1 #3 | ACAAAAATATTGTCAAATT |
| siIRF3 #1 | CGGAGCAAGGACCCTCACG |
| siIRF3 #2 | AAGAGGCTCGTGATGGTCA |
| siIRF3 #3 | CGGAAGGAAGCGGACGCTC |
| siIRF3 #4 | CCCTTCATTGTAGATCTGA |
| sip65 #1 | CCTTCCAAGTTCCTATAGA |
| sip65 #2 | AAGCTGATGTGCACCGACA |
| Real-time qPCR | |
| CCL2 forward primer | AGAATCACCAGCAGCAAGTGTCC |
| CCL2 reverse primer | TTGCTTGTCCAGGTGGTCCATG |
| CCL5 forward primer | CCTCGCTGTCATCCTCATTGCTAC |
| CCL5 reverse primer | CTTGACCTGTGGACGACTGCTG |

| | |
|---|---|
| CCL20 forward primer | GCTCCTGGCTGCTTTGATGT |
| CCL20 reverse primer | GCCGTGTGAAGCCCACAATA |
| CXCL10 forward primer | CTGCCATTCTGATTTGCTGCCTTATC |
| CXCL10 reverse primer | GATGCAGGTACAGCGTACAGTTCTAG |
| β-actin forward primer | CGGGAAATCGTGCGTGAC |
| β-actin reverse primer | TGGAAGGTGGACAGCGAGG |

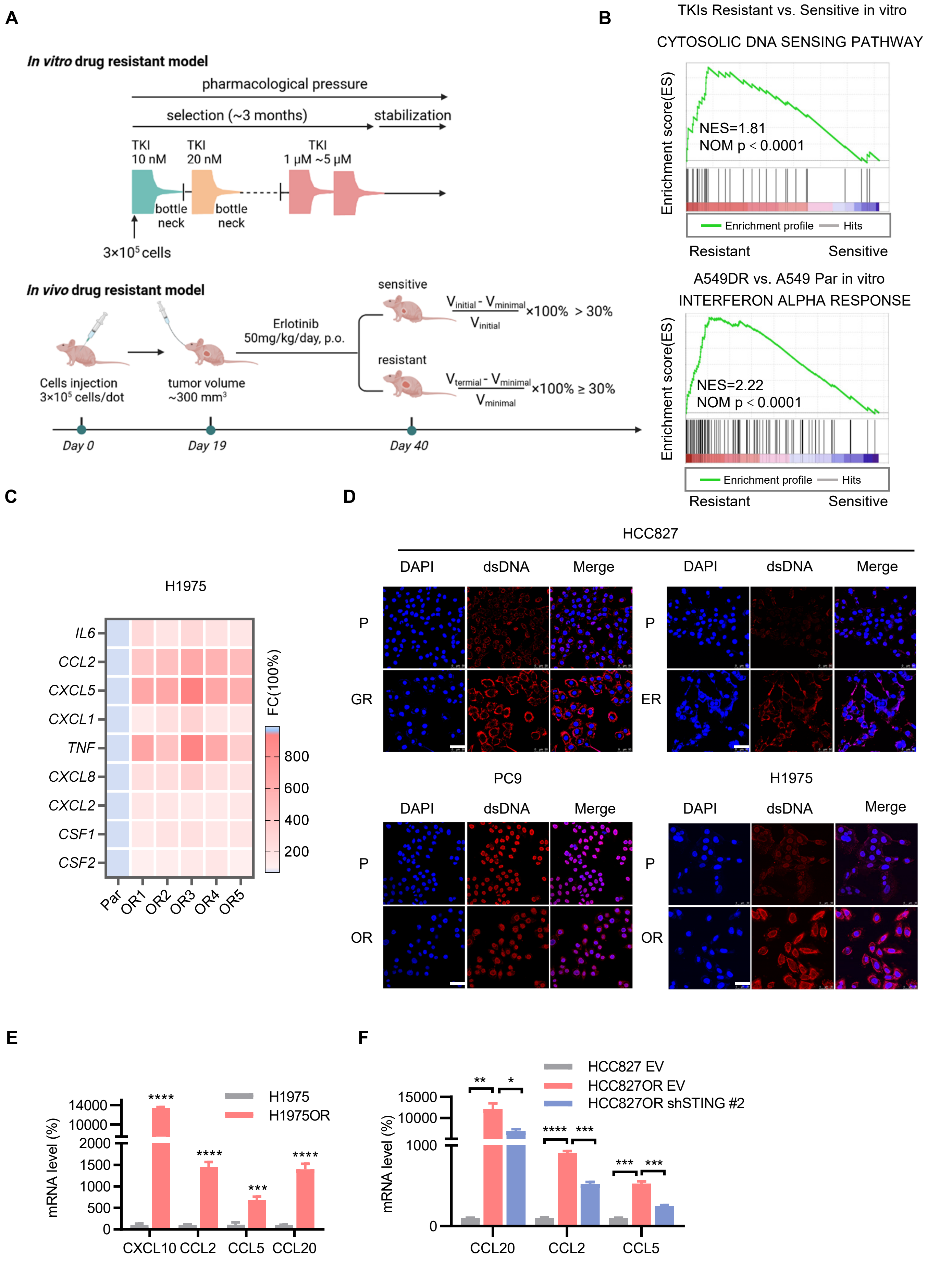


$$\frac{V_{initial} - V_{minimal}}{V_{initial}} \times 100\% > 30\%$$

$$\frac{V_{termial} - V_{minimal}}{V_{minimal}} \times 100\% \geq 30\%$$

**Supplementary Fig. S1**

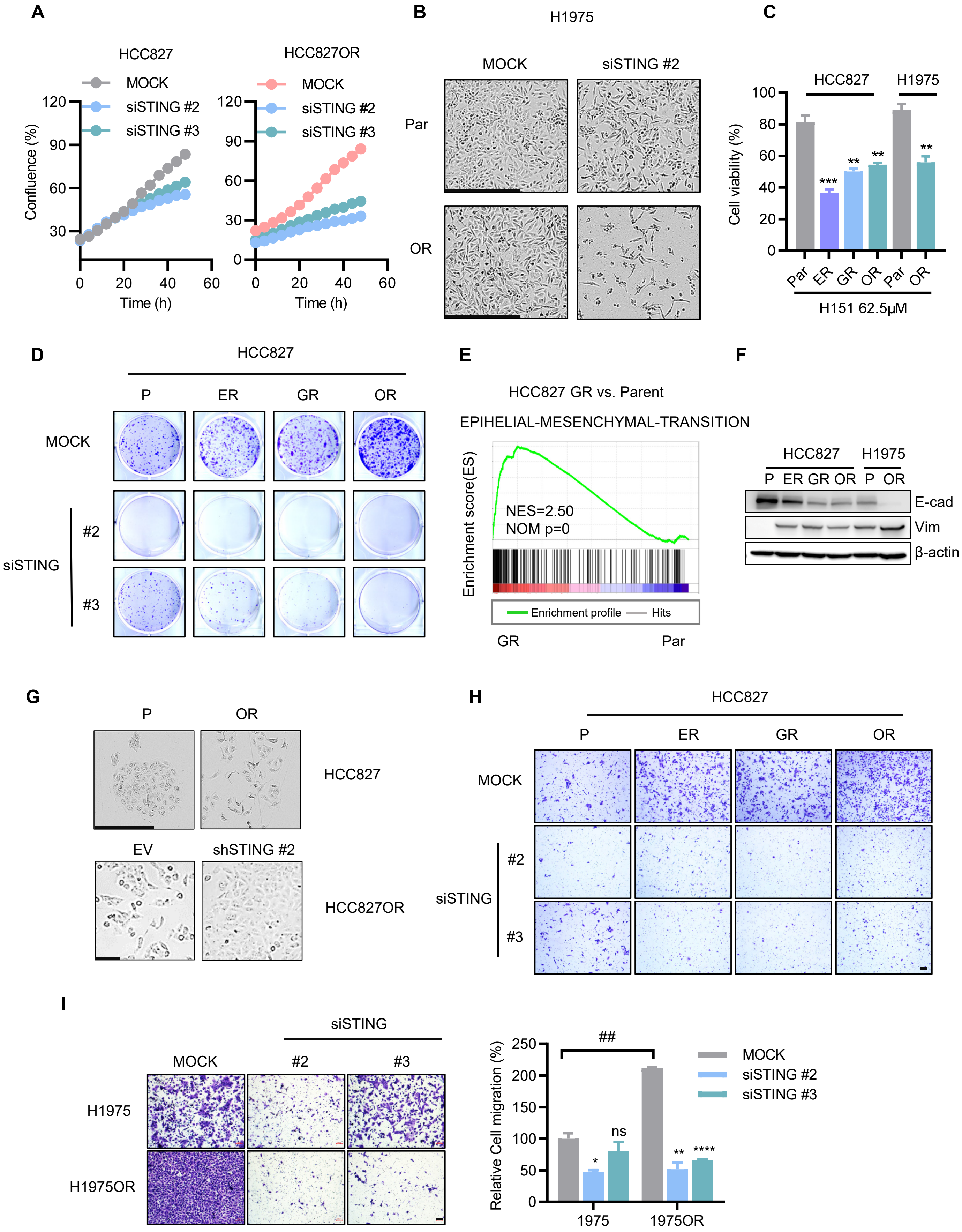


**Supplementary Fig. S2**

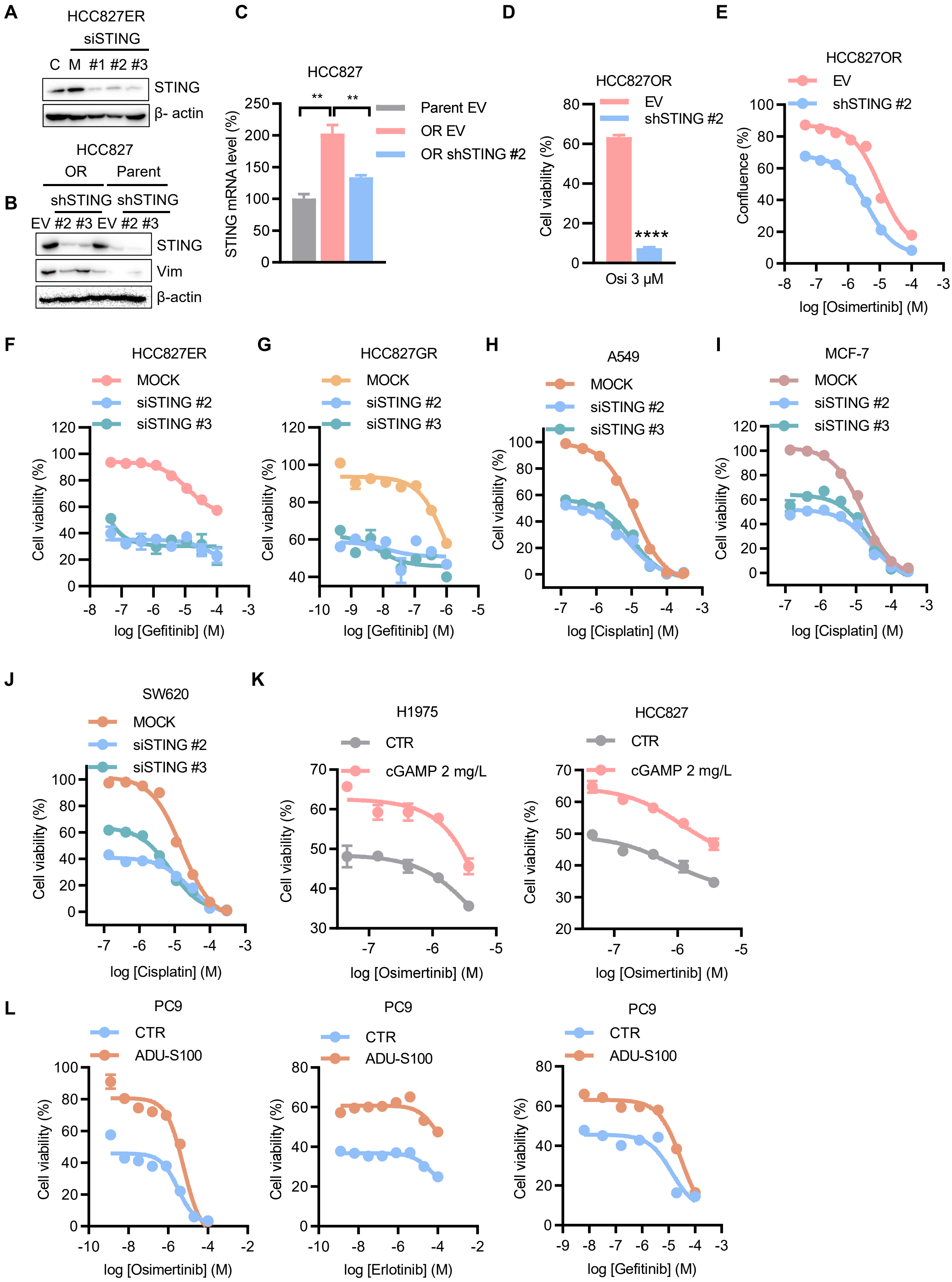


**Supplementary Fig. S3**

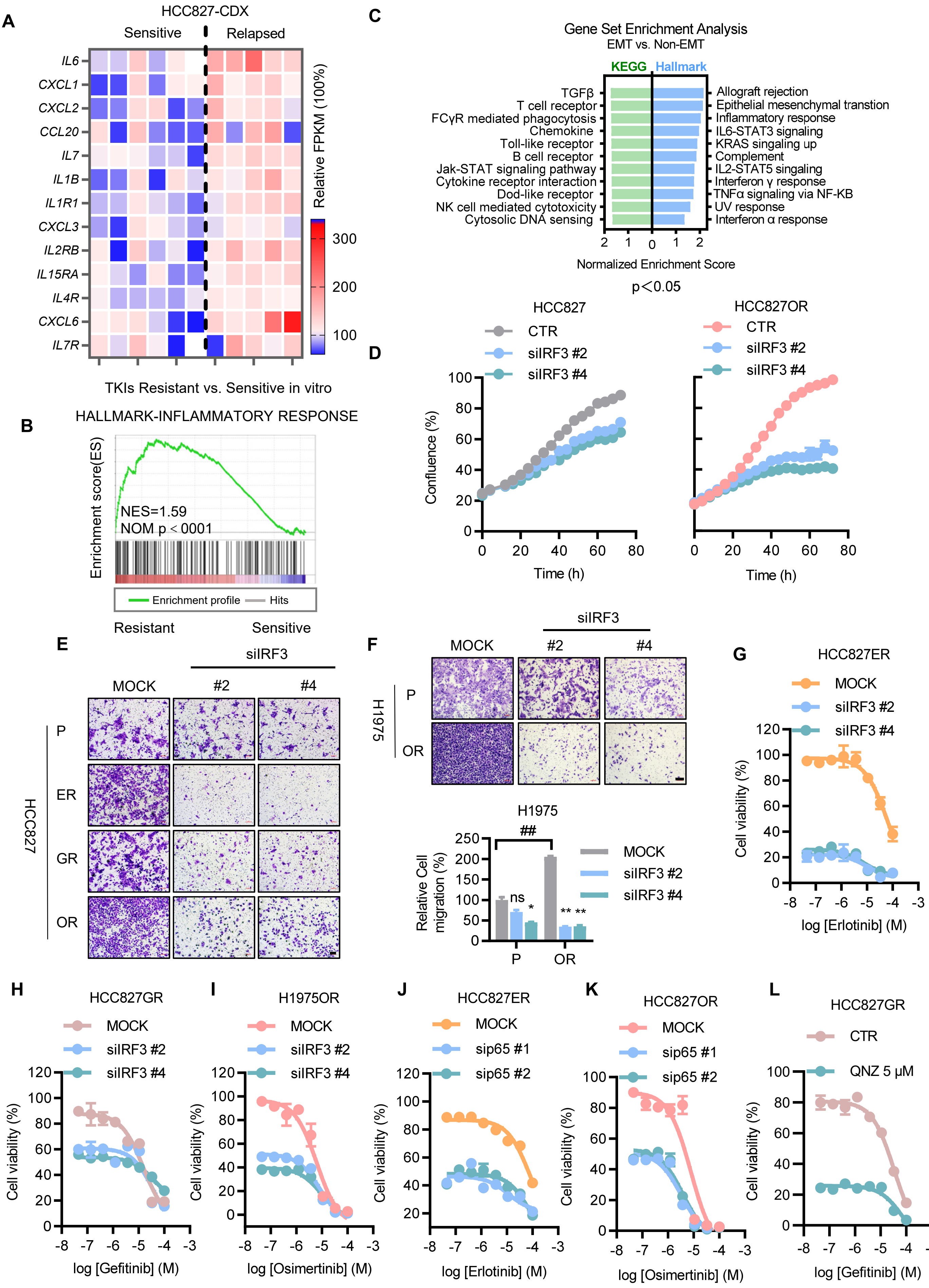


**Supplementary Fig. S4**

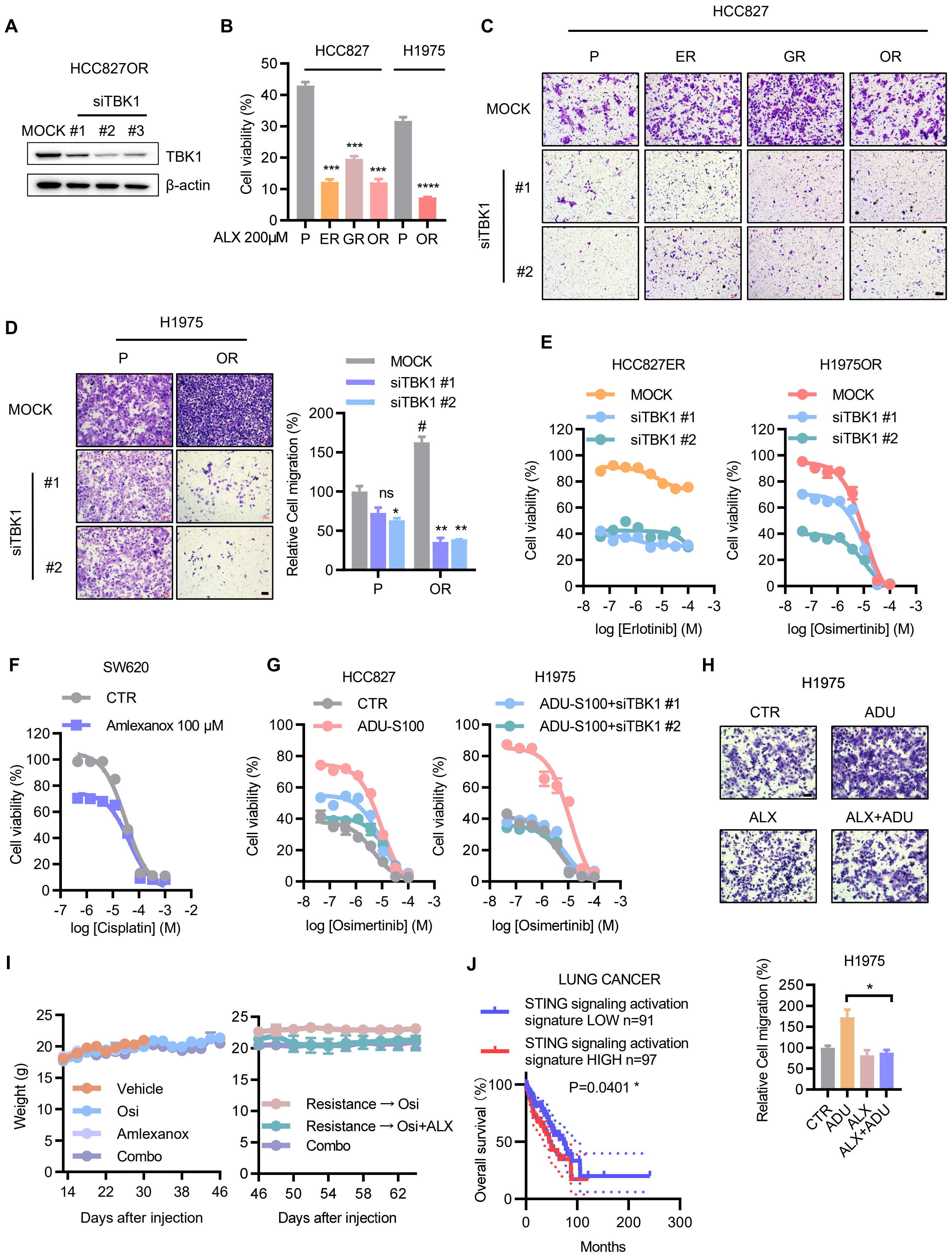


**Supplementary Fig. S5**
